# Glucose Uptake as an Alternative to Stop-Flow Respirometry for Measuring Metabolic Rate in *Danio rerio* Larvae

**DOI:** 10.1101/2021.04.30.442098

**Authors:** Bridget L Evans, Adam F L Hurlstone, Peter E Clayton, Adam Stevens, Holly A Shiels

## Abstract

Respirometry is the current gold-standard for measuring metabolic rate. However, there is a growing need for metabolic rate measurements suitable for developmental studies, particularly in *Danio rerio*, where many important developmental stages occur at < 4 mm. While many metabolic studies rely on respirometry, the cost and complexity of the equipment limits its appeal in non-specialist labs, and background respiration becomes increasingly problematic as the size of the organism reduces. Here, glucose uptake was compared to stop-flow respirometry as an alternative measure of metabolic rate more suitable to the small scale required for developmental studies. A Passing-Bablok regression revealed the rate of glucose uptake can be considered equivalent to oxygen consumption as a measure of metabolic rate in *Danio rerio* larvae within a 95% limit of agreement. Therefore, glucose uptake is a valid alternative to the gold-standard in small organisms where conventional respirometry is problematic.

**Summary statement:** The rate of glucose uptake is a valid alternative to respirometry for metabolic rate measurements in small larval fish.

## Introduction

Metabolism encompasses a complex network of chemical reactions fundamental for sustaining life. Metabolic reactions convert chemicals from one form to another to release energy (e.g., respiration), generate molecules required for other processes (such as amino acids and nucleotides), and facilitate waste removal (including CO_2_ and nitrogen).

Due to the complicated nature of whole-organism metabolism, reductionist approaches are often used to identify and measure single, key metabolic pathways (Darden, 2016). This enables the standardisation of metabolic rate measurements and facilitates comparisons between different studies. Pathways in the metabolic network are generally targeted for measurement under the assumption that a complex system is the sum of its parts, and thus each step is carried out in proportion to any other step, and that any limiting factor will limit the entire system as opposed to an individual step.

Key steps that have been targeted in metabolic research include the rate of substrate consumption, rate of generation and excretion of metabolites, and production of biproducts. For example, respirometry systems are employed to measure the rate of oxygen consumption at the level of the whole organism, the tissue, or the isolated mitochondria. Respirometry may also be used to measure production of carbon dioxide (Tickle, Hutchinson and Codd, 2018), while metabolite production can be detected in cells using analysers such as the Agilent Seahorse (Agilent, Santa Clara, California, US)(Müller *et al*., 2019). Glucose tolerance testing is used to measure glucose uptake, while calorimetry, one of the oldest methods of estimating metabolic rate, measures heat production, a biproduct of many chemical reactions (Gillis *et al*., 2015).

One of the most important metabolic pathways in biological systems is glucose metabolism, the mechanism by which glucose is processed to generate ATP, the primary unit of energy in living systems. Glucose is uptaken by respiring cells via glucose transporters and immediately phosphorylated to glucose-6-phosphate (G6P) by hexokinases. This prevents diffusion out of the cell, as G6P is membrane impermeable, accumulating G6P as a substrate for energy production. G6P is modified in multiple sequential reactions, ultimately generating one pyruvate molecule, and releasing 2 ATP per glucose molecule via glycolysis. This is the final step in anaerobic respiration. Pyruvate is subsequently transported from the cytosol into the mitochondria to enter the citric acid cycle and commence aerobic respiration. NADH generated by the citric acid cycle fuels the electron transport chain, ultimately generating a further 36 ATP. Glucose and oxygen are both limiting factors in glucose metabolism:

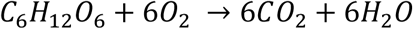

and can therefore act as indicators of metabolic rate.

While respirometry is considered the gold-standard for measuring metabolism, respirometry systems are often complex (see figure 1) and depending on the level of biological organization under investigation can require significant setup and validation time. They can also be costly, needing specialist equipment, particular for micro-respirometry (such as the Oroboros Oxygraph-2k and Agilent Seahorse XF Analyzer). While whole animal respirometry systems are less expensive, they require specialist knowledge to ensure reliability of experimental design and interpretation (Clark, Sandblom and Jutfelt, 2013). Common errors associated with whole animal measurements are compounded when considering embryos and larvae. Background bacterial respiration represents one of the greatest sources of interference in respirometry systems (Clark, Sandblom and Jutfelt, 2013)(M. B. S. Svendsen, Bushnell and Steffensen, 2016). While this can be negligible when compared with respiration of larger organisms, the interference increases as the size of the organism decreases (M. B.S. Svendsen, Bushnell and Steffensen, 2016). While reducing the volume of the respirometry setup limits the impact of bacterial respiration, the volume and capacity of the system can only be reduced a finite amount and often requires more specialist, expensive equipment.

**Figure 1:**
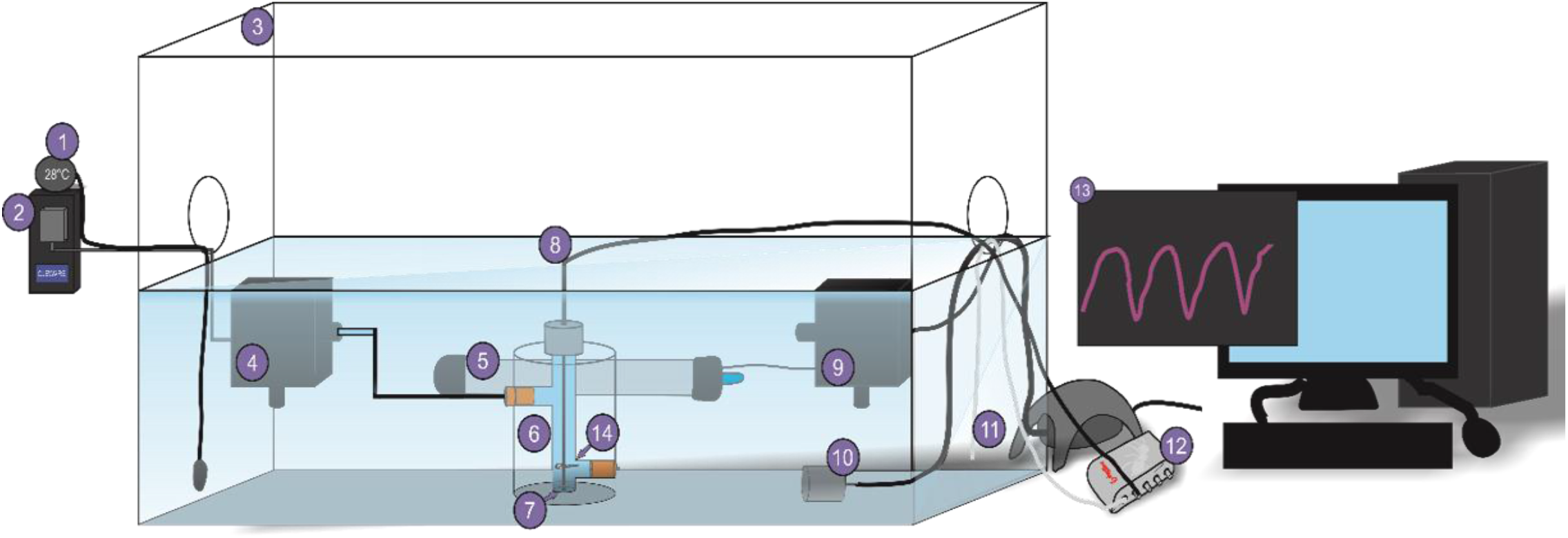
Stop-flow respirometry setup for larval fish. ***1***. Temperature probe. ***2***. AquaResp-controlled USB switch. ***3***. Recirculation tank. ***4***. Stop-flow pump controlled by USB switch. ***5***. Water heater (28°C). ***6***. Respirometry chamber. ***7***. Oxygen spot. ***8***. Fibreoptic probe. ***9***. Recirculation pump. ***10***. Air stone. ***11***. Temperature probe. ***12***. FireStingO_2_ Oxygen and Temperature Logger. ***13***. AquaResp software v.3. Larvae (***14***.) are placed in the respirometry chamber and acclimated for one hour before measurements begin. AquaResp controls the stop-flow pump and logger to cycle through a series of flush-wait-measure steps (60 s, 10 s, 300 s), generating oxygen saturation curves. AquaResp generates regression curves and outputs MO_2_ data for each cycle.

Glucose uptake assays, on the other hand, do not require specialist equipment to perform: only a plate reader, microscope, and reagents are needed. These assays can be adapted to 96 well plate formats, enabling relatively high throughput, and higher granularity. A glucose uptake assay measures the rate of uptake of a glucose analogue, 2-deoxyglucose (2DG), which is transported into the cell by glucose transporters. Hexokinases phosphorylate 2DG into 2-deoxyglucose-6-phosphate (2DG6P), which is membrane impermeable and resistant to further modification, subsequently accumulating in the cell at the same rate as glucose uptake. Glucose-6-phosphate dehydrogenase converts 2DG6P into 6-phosphodeoxygluconate, reducing NADP^+^ into NADPH. Reductases catabolise the conversion of proluciferin to luciferin, oxidising NADPH to NADP^+^, producing luminescence proportional to the rate of glucose uptake (see *Figure 2a*). Unlike respirometry systems, glucose uptake assays do not require specialist software to interpret and process data. The raw luminescence data can be used directly without the need for complicated respirometry calculations. Set-up time is minimal, as the assay can be performed as soon as the reagents arrive, and the cost is comparatively inexpensive compared to purchasing a respirometry setup.

**Figure 2:**
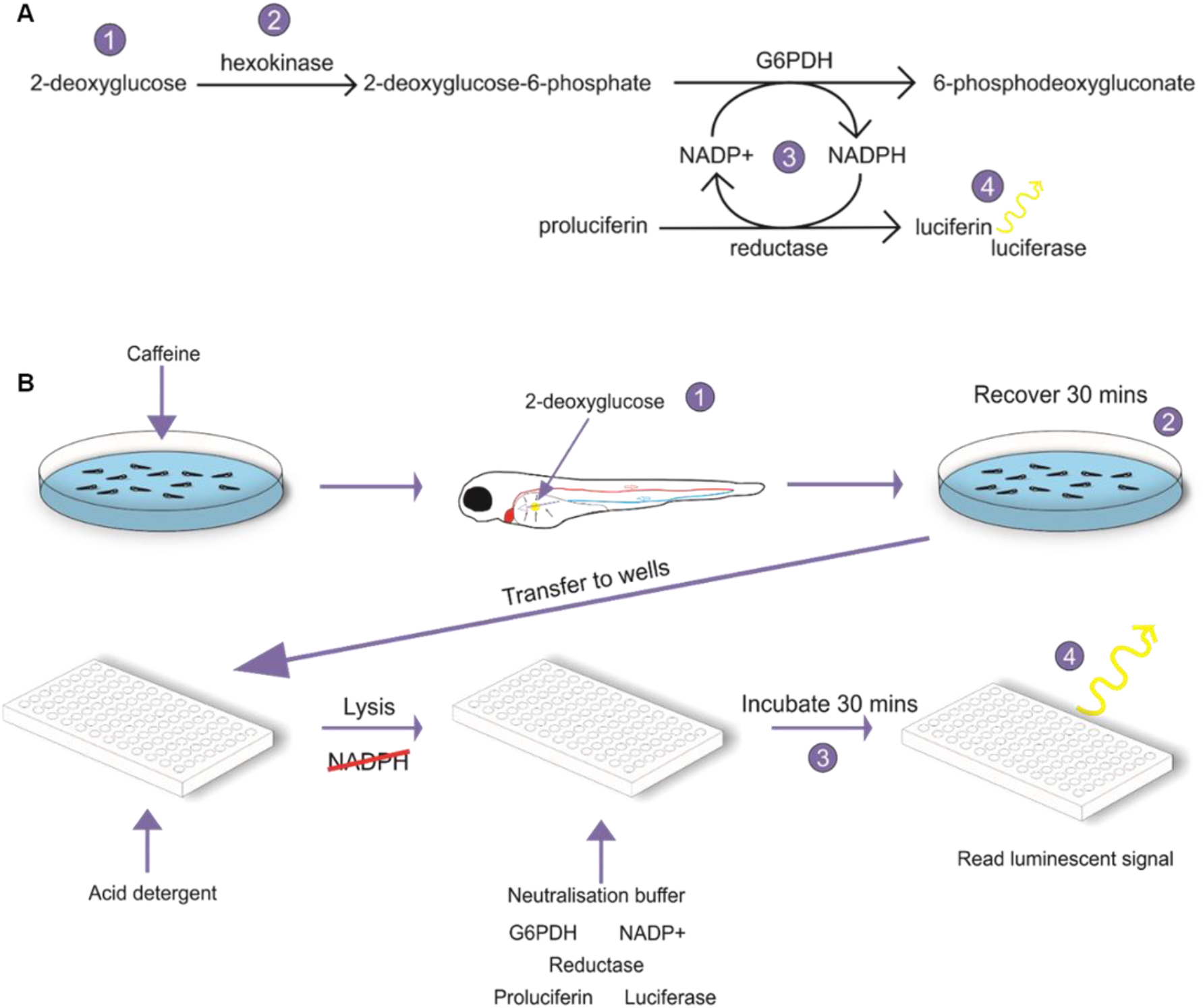
Glucose uptake assay protocol. ***2a***. Equation detailing the reactions involved in a glucose uptake assay. ***2b***. Experimental setup of a glucose assay performed on 96 hpf zebrafish larvae exposed to different concentrations of caffeine. Larvae are injected with 2-deoxyglucose into the yolk (yellow area) and recovered for 30 minutes. Larvae are transferred to individual wells and lysed with acid detergent to stop further glucose uptake, destroy any native NADPH, and homogenise the sample. Neutralisation and detection buffer are added, and samples are incubated while the luminescent signal is generated. After 30 minutes, luminescence is measured on a plate reader.

Zebrafish (*Danio rerio*) are increasingly used in metabolic studies, and are an invaluable model organism for developmental research, featuring a rapid generation time, large spawn size, genetic tractability, and transparent embryos (Howe *et al*., 2013). There is a growing body of literature incorporating metabolism into developmental studies in zebrafish (Dhillon *et al*., 2019)(Roy *et al*., 2017)(Seth, Stemple and Barroso, 2013), and several important developmental stages occur in the zebrafish before 4 mm total body length, so a method for reliably measuring metabolism at smaller sizes would be beneficial. In zebrafish larvae the yolk serves as a primary energy store. The developing circulatory system runs directly across the yolk, into the heart, dorsal aorta, caudal vein, and back to the yolk in one continuous circuit. Nutrients dissolve into the bloodstream when passing through the yolk, which are then absorbed by actively respiring cells. As the yolk is readily accessible, a glucose analogue can be introduced easily into the system, provide the means to assess rate of glucose uptake as an index of metabolic rate.

Here, a glucose uptake assay in 96 hours post-fertilisation (hpf) zebrafish larva, adapted from an existing cell-culture protocol, is compared with whole-organism metabolic rate measured using respirometry. Protocol comparison shows the two methods are equivalent, thus the glucose uptake assay can be used as a new proxy for whole animal metabolism in larval fish.

## Materials and Methods

### Zebrafish Husbandry

An established line of AB Notts zebrafish from the Biological Services Unit of the University of Manchester were used in this study. Adult zebrafish were housed under standard conditions (≈ 28°C; 14 h light/10 h dark cycle; stocking density < 5 fish per litre). Breeding pairs were separated and housed in breeding tanks overnight at a ratio of one male to one female. Dividers were removed at the start of the following light cycle. Embryos were collected after one hour of free breeding and incubated in embryo water (Instant Ocean salt 60 µgml^-1^, 2 µgml^-1^ methylene blue (Dunn, 2018)) at 28°C at a stocking density of < 50 embryos per petri dish. Unfertilised embryos were removed after 24 hours and embryo water was refreshed every 24 hours.

### Caffeine Treatment

Caffeine is a well-known stimulant inducing multiple effects on the organism, including increased heart rate, metabolic rate, and locomotion (Rana *et al*., 2010)(Abdelkader *et al*., 2012)(Santos *et al*., 2017)(Chen *et al*., 2008). To generate embryos with differing metabolic rates, varying concentrations of caffeine were dissolved in embryo water (0 mgL^-1^, 5 mgL^-1^, and 25 mgL^-1^). Concentrations were selected to reliably induce a metabolic effect (Rana *et al*., 2010)(Abdelkader *et al*., 2012)(Santos *et al*., 2017)(Chen *et al*., 2008). Prior to measuring indices of metabolism, larvae were randomly assigned to petri dishes containing one of the three concentrations of caffeine and returned to the 28°C incubator for two hours. Larvae were subsequently assigned to either stop-flow respirometry or the glucose uptake assay. Age in hpf has a notable impact on metabolic rate (Dhillon *et al*., 2019), so experiments were performed on age matched separate individuals.

### Oxygen Consumption via Closed-Circuit Stop-Flow Respirometry

In order to accurately measure very small changes in oxygen saturation, a closed-circuit stop-flow respirometer was designed (*Figure 1*) (M. B.S. Svendsen, Bushnell and Steffensen, 2016), with the volume of the respirometer kept to a minimum (3 ml). Flo-thru probe vessels (*Figure 1.6*) (Loligo Systems, 3-6 mm probe/10 mm tube size) were adapted for use as respirometry chambers. Single oxygen spots (*Figure 1.7*) (Pyroscience, Aachen, Germany) were placed at the base of each chamber, paired with optical oxygen sensors (*Figure 1.8*) (Pyroscience, Aachen, Germany) fed through the top of the chamber, calibrated according to the manufacturer’s instructions. A stop-flow pump was attached to each chamber (*Figure 1.4*), controlled by a Cleware USB-Switch (*Figure 1.2*) (Cleware GmbH, Germany) programmable switch and AquaResp v.3 software (*Figure 1.13*) (AquaResp, v3, Python 3.6, (Svendsen, Bushnell and Steffensen, 2019)). Cycling parameters were set to 60 s, 10 s, 300 s, (flush, wait, measure). This was sufficient flush time to replace all the water within the chamber with oxygenated water, and a sufficient wait time to ensure complete mixing within the chamber. To maintain a constant temperature of 28°C ± 0.3°C (*Figure 1.1*), the respirometry system was immersed in a recirculation chamber (*Figure 1.3*) under constant aeration (*Figure 1.10*). During flushing, water was pumped from this water bath into the chambers. Oxygen saturation within the chambers and water temperature (*Figure 1.11*) was recorded using a FireStingO_2_ Fiber-optic oxygen and temperature meter (*Figure 1.12*) (PyroScience GmbH, Aachen, Germany) and Pyroscience Pyro Oxygen Logger (PyroScience GmbH, Aachen, Germany, v3.312, firmware 3.07).

Prior to each trial, the respirometer was filled with water matching the treatment group. Individual zebrafish larvae (96 hpf) were placed into a random chamber and allowed to acclimate for 30 minutes prior to the first trial. Data are represented as the mean of three trials per larvae (n = 10 per treatment). Oxygen consumption was measured on empty chambers immediately before and after each set of trials to account for background bacterial respiration.

Regression curves were automatically generated by AquaResp and oxygen consumption was extracted and normalised against the length of the trial and volume of the respirometry chamber according to the following equation, where 7.8 mgL^-1^ is the maximum dissolved oxygen in 28°C desalinated water:

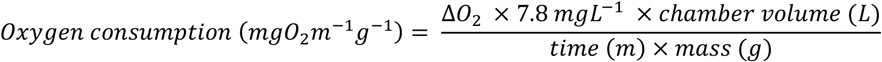

### Glucose Uptake Assay

An adapted Glucose Uptake-Glo™ Assay (Promega, Wisconsin, USA) was performed on 96 hpf zebrafish to assess differences in the rate of glucose uptake between treatment groups. Following the incubation in caffeine solution, larvae were immobilised in 0.7 µM MS-222 (no impact on heart rate (Zakaria *et al*., 2018)) and methylcellulose. Individual zebrafish larvae (n = 10 per treatment) were injected with 1 mM 2DG (*Figure 2.1*) directly into the yolk and recovered in caffeinated water for 30 minutes (*Figure 2.2*), allowing the 2DG to disperse into the yolk and be transported into the cells via the bloodstream. Larvae were subsequently terminated in MS-222 (500 mgL^-1^), transferred to a 96 well plate, and homogenised in acid detergent with a fine gauge needle to terminate glucose uptake and destroy NADPH present in the sample. Each well was neutralised with a high pH buffer and incubated with detection reagent (G6PDH, NADP^+^, reductase, UltraGlo™ recombinant luciferase, proluciferin) for 30 minutes. G6PDH oxidises accumulated 2DG6P to 6-phosphodeoxygluconate, reducing NADP^+^ to NADPH. Reductase catalyses the production of luciferin from proluciferin, oxidising NADPH to NADP^+^ (*Figure 2.3*). Luciferin acts as a substrate for the UltraGlo™ recombinant luciferase to generate luminescence (*Figure 2.4*). As all native NADPH is removed from the sample during the lysis step, luminescence ↔ NADPH ↔ 2DG6P accumulation ↔ glucose uptake. The luminescent signal is therefore proportional to the rate of glucose uptake. To limit the impact of native glucose interference on the signal, sham-injected larvae were included as a control to adjust for background luminescence. Sham-injected larvae received an injection of phenol red, a common innocuous tracer used in microinjection solutions, in place of the 2DG. An empty well was included per row to further minimise background noise. Luminescence was measured on a Synergy™ H1 Microplate reader (BioTek Instruments, Inc., Winooski, VT, USA, software version 2.07.17) with 8 readings per well, adjusted against the background signal.

### Statistical Tests

Data were imported into GraphPad Prism (v7.04) for statistical analysis and converted to logarithmic values. All data were ROUT tested for outliers (Motulsky and Brown, 2006) and subject to D’Agostino and Pearson normality tests. Post-hoc power calculations were performed to confirm sample sizes were sufficient, where α = 0.05. One-way analysis of variance (ANOVA) was used to determine if there was a significant difference between the caffeine treatments within a given metabolic index.

To compare between the two methods of assessing metabolism, Passing-Bablok regression analysis was used, where a regression line is fitted for the alternative method against the gold-standard (Bablok, 1983; Bilic-Zulle, 2011). The Passing-Bablok regression requires the data to be linear, and states that if a structural relationship exists between two methods, it can be described by the linear equation *y = α + βx*. If the 95% confidence intervals of *α* include 0 and *β* include 1, the two methods are comparable within the given range. If the confidence interval of *α* does not include 0, there is a systematic difference between the methods, and if the confidence interval of *β* does not include 1, there is a proportional difference between the methods. The Passing-Bablok regression 95% confidence bounds were calculated using the jack-knife method. The regression plot is presented alongside a Bland-Altman plot of the residuals, together with the bias and limits of agreement (Giavarina, 2015). The bias 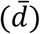 was calculated as the mean of the differences between the two methods. The limits of agreement were calculated as:

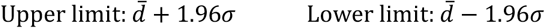

where n is the number of individuals (30) and σ is the standard deviation. The upper and lower 95% confidence intervals were calculated as:

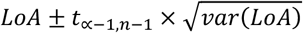

Where:

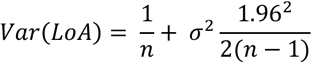

The residuals were plotted as the percentage difference [(Respirometry – Glucose Uptake)/(Respirometry)*100] vs the averages to account for the increase in variability as the magnitude of the measurement increased (Giavarina, 2015).

To calculate the biological power, the literature was reviewed to ascertain normal variation between individual zebrafish larvae and the expected percentage difference between groups with different metabolic rates. PASS 2021 (Power Analysis & Sample Size, NCSS Statistical Software, Utah, USA, v21.0.2) (Bland and Altman, 2010; Lu *et al*., 2016) was used to calculate the power associated with the defined maximum allowable difference at a confidence level of 85%.

## Results and Discussion

Metabolic rate measured by whole animal oxygen consumption and by glucose uptake were elevated with increasing caffeine concentration (*Figure 3*). Oxygen consumption sequentially and significantly increased between control, 5 mgL^-1^, and 25 mgL^-1^ (*Figure 3a*), reflecting the increase in movement, heart rate, and metabolic demand induced by caffeine exposure. Oxygen is required for aerobic respiration and can be considered a limiting factor of respiratory rate. Therefore, the increase in oxygen consumption observed with caffeine exposure in 96hpf zebrafish larva can be considered a reliable measure of increasing organismal metabolic rate.

**Figure 3:**
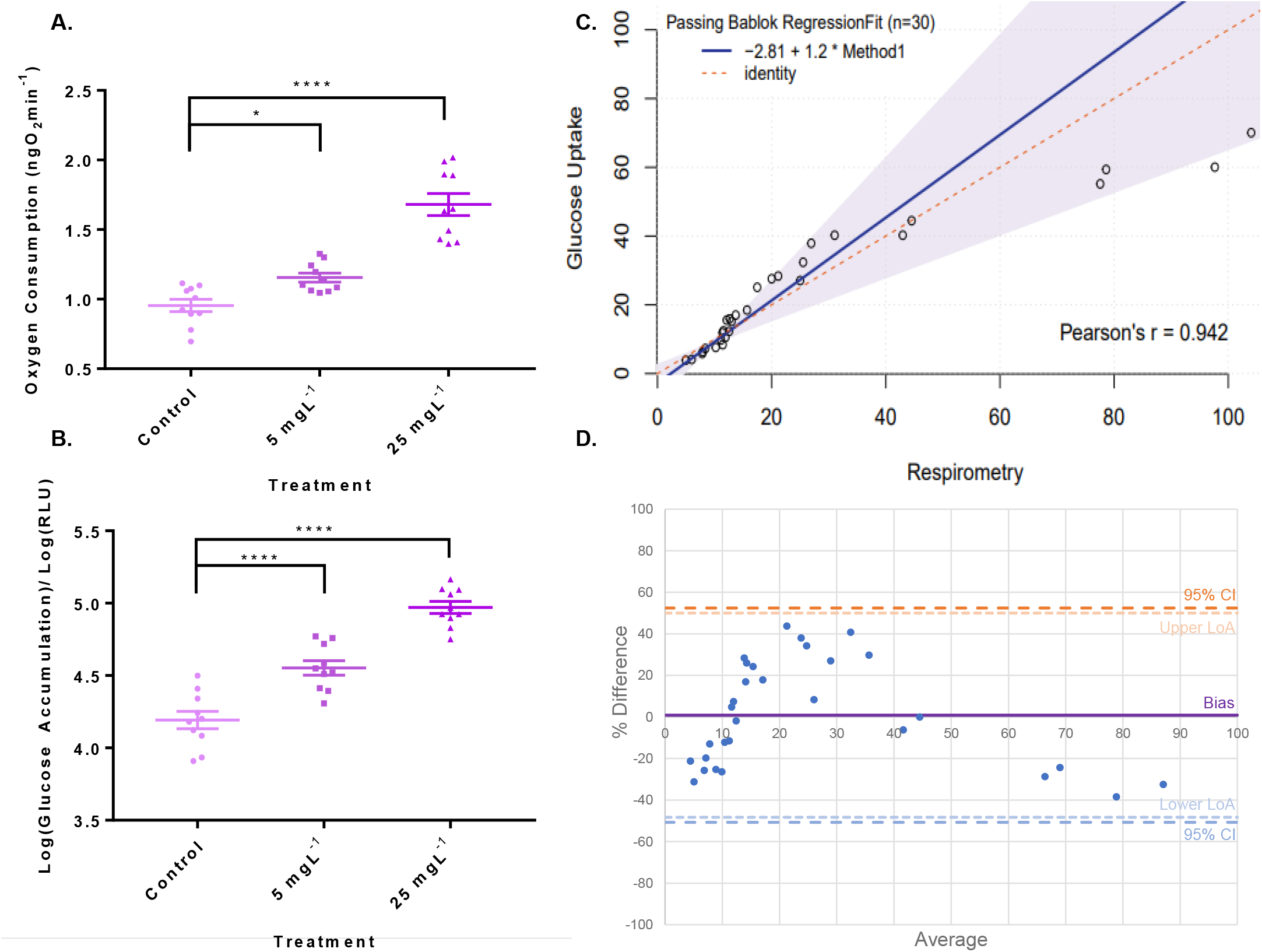
Comparison of metabolic rate as measured by stop-flow respirometry (3a) and glucose uptake (3b), Passing-Bablok comparison of the two methods (3c), and Bland-Altman residuals plot (3d). 96 hpf zebrafish larvae were incubated in one of three caffeine concentrations (0 mgL^-1^, 5 mgL^-1^, and 25 mgL^-1^) to generate groups with differing metabolic rates. ***3a***. Oxygen consumption was measured by stop-flow respirometry in milligrams of oxygen per minute per fish (n = 10). There was a significant increase in oxygen consumption between control and caffeine treated zebrafish larvae, with oxygen consumption increasing with increasing concentrations of caffeine (* = 0.0429. **** <0.0001). ***3b***. The rate of glucose uptake was measured using a Glucose-Uptake Glo™ Assay. There was a significant increase in luminescence between control and caffeine treated zebrafish larvae, with higher levels of luminescence correlating with higher concentrations of caffeine (**** <0.0001). All data are represented as individual data points with mean and SEM bars. Significance was determined by one-way ANOVA. ***3c***. Passing-Bablok Regression of respirometry vs. scaled glucose assay data, including regression line and 95% confidence limits. *y = 2.81 + 1.20x*, 95% confidence intervals -8.50 ≤ *α* ≤ 2.87 and 0.28 ≤ *β* ≤1.79. ***3d***. Bland-Altman plot of residuals of the log values. Bias = 0.85%, upper limit of agreement = 49.99%, lower limit of agreement = -48.33%, upper confidence interval = 52.43%, lower confidence interval = -50.74%.

Similarly, glucose accumulation in zebrafish larvae significantly increased as caffeine concentration increased (*Figure 3b*). As glucose is the initial substrate of glycolytic respiration, glucose is also a limiting factor of metabolism. Therefore, akin to changes in oxygen, glucose accumulation can be considered a measure of whole animal metabolic rate. While aerobic respiration generates the majority of the ATP produced during glucose metabolism (36 units ATP per glucose, 6 units ATP per oxygen), anaerobic respiration also contributes to overall energy production (2 units ATP per glucose [anaerobic] vs 36 units ATP per glucose [aerobic]). Stop-flow respirometry measures changes in oxygen consumption and can thus only serve as an indicator of aerobic metabolism. Glucose accumulation measures the limiting factor earlier in the pathway and thus can account for both aerobic and anaerobic metabolism. The differences in percentage change between treatment groups observed in the two methods is likely due to this difference in scope of the two methods: aerobic (respirometry) and aerobic + anaerobic (glucose accumulation).

The Passing-Bablok regression was used to compare and assess the equivalence of the two methods. D’Agostino and Pearson normality testing of each data group showed data were from a normal distribution (p > 0.1). Regression of the raw data (*Figure 3c*) gave the following regression line:

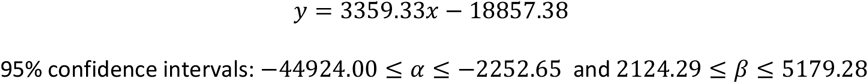

As the 95% confidence intervals for α do not include 0 and the intervals for β do not include 1, there is a proportional and systematic difference between the two methods. This is expected, as the two methods use different units, and can be corrected for by scaling the arbitrary glucose accumulation data by 0.00048. The parameters of the scaled data for the Passing-Bablok regression were:

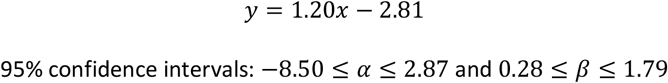

As both *α* = 0 and *β* = 1 are true at a confidence of 95%, it can be concluded that the two methods are comparable and interchangeable.

The residuals were plotted as percentage difference vs average of the two methods alongside the bias (0.85% {-8.52, 10.21}) and the upper (49.99%) and lower (−48.30%) limits of agreement in a Bland-Altman plot (*Figure 3d*). The 95% confidence interval of the bias included zero, indicating there is no statistically significant bias, confirmed by a two-tailed t-test (p = 0.893). As shown in *Figure 3d*, 100% of the data fell between the upper and lower limits of agreement, suggesting the residuals are distributed around the bias line. Finally, existing literature was used to assess the biological power of the comparison. As metabolic rate can vary between zebrafish embryos by up to approximately 50% (Barrionuevo and Burggren, 1999; Bang, Gronkjaer and Malte, 2004), a difference of this size can be considered within normal metabolic variation and not biologically significant. As shown in *Figure 3d*, the data falls within the 95% limits of agreement, suggesting the differences between these two methods are not likely to be biologically relevant. To determine the power of the method, a further 10% difference in metabolic rate was set as the minimum requirement for biological significance (60% total). The power was subsequently calculated using the approximate maximum allowable difference of 59%. This study achieves a power of 94.99%, based on a mean difference of 0.85% and a standard deviation of 25.07%, suggesting the difference between the two methods is unlikely to mask biologically relevant differences between groups, and the methods can likely be used interchangeably.

In summary, regression analysis shows that glucose accumulation and oxygen consumption measurements can be considered equivalent when measuring differences in metabolic rate induced by caffeine exposure in larval zebrafish. While respirometry yields information on aerobic metabolic rate and has been readily used on a variety of organisms, the practical and financial aspects of setting up and validating a respirometry system reduces their appeal, particularly for very small animals. Glucose accumulation, though limited to smaller organisms, represents a readily accessible, rapid, and simple alternative to stop-flow respirometry in larval fish species.

## Abbreviations

2DG: 2-Deoxyglucose
2DG6P: 2-Deoxyglucose-6-Phosphate
ANOVA: Analysis of Variance
ATP: Adenosine Triphosphate
G6P: Glucose-6-Phosphate
G6PDH: Glucose-6-Phosphate Dehydrogenase
MO_2_: Oxygen Consumption Rate
MS-222: Tricaine Methanesulfonate
SEM: Standard Error of Mean

## Acknowledgements

The authors would also like to acknowledge the Biological Services Unit at the University of Manchester for housing and maintaining the zebrafish lines.

## Author contributions

**Bridget Evans:** Conceptualisation, methodology, validation, formal analysis, investigation, writing – original draft preparation, review, and editing, and visualisation.

**Holly Shiels:** Conceptualisation, writing – review and editing, supervision.

**Adam Hurlstone:** Conceptualisation, resources, writing – review and editing, supervision.

**Adam Stevens:** writing – review, supervision.

**Peter Clayton:** writing – review, supervision.

## References

1. Abdelkader, T. S. et al.. (2012) ‘Exposure time to caffeine affects heartbeat and cell damage-related gene expression of zebrafish Danio rerio embryos at early developmental stages’, Journal of Applied Toxicology. John Wiley & Sons, Ltd, 33(11), p. n/a-n/a. doi: 10.1002/jat.2787.

2. Bablok, W. (1983) ‘Ein neues biometrisches Verfahren zur Überprüfung der Gleichheit von Meßwerten von zwei analytischen Methoden: Anwendung von linearen Regressionsverfahren bei Methodenvergleichsstudien in der Klinischen Chemie, Teil I’, Clinical Chemistry and Laboratory Medicine. Walter de Gruyter, Berlin / New York, 21(11), pp. 709–720. doi: 10.1515/cclm.1983.21.11.709.

3. Bang, A., Gronkjaer, P. and Malte, H. (2004) ‘Individual variation in the rate of oxygen consumption by zebrafish embryos’, Journal of Fish Biology. John Wiley & Sons, Ltd, 64(5), pp. 1285–1296. doi: 10.1111/j.0022-1112.2004.00391.x.

4. Barrionuevo, W. R. and Burggren, W. W. (1999) O 2 consumption and heart rate in developing zebrafish (Danio rerio): influence of temperature and ambient O 2.

5. Bilic-Zulle, L. (2011) ‘Comparison of methods: Passing and Bablok regression’, Biochemia Medica. Biochemia Medica, Editorial Office, 21(1), pp. 49–52. doi: 10.11613/BM.2011.010.

6. Bland, J. M. and Altman, D. G. (2010) ‘Statistical methods for assessing agreement between two methods of clinical measurement’, International Journal of Nursing Studies, pp. 931–936. doi: 10.1016/j.ijnurstu.2009.10.001.

7. Chen, Y. H. et al.. (2008) ‘Movement disorder and neuromuscular change in zebrafish embryos after exposure to caffeine’, Neurotoxicology and Teratology. Neurotoxicol Teratol, 30(5), pp. 440–447. doi: 10.1016/j.ntt.2008.04.003.

8. Clark, T. D., Sandblom, E. and Jutfelt, F. (2013) ‘Aerobic scope measurements of fishes in an era of climate change: Respirometry, relevance and recommendations’, Journal of Experimental Biology. J Exp Biol, pp. 2771–2782. doi: 10.1242/jeb.084251.

9. Darden, L. (2016) ‘Reductionism in Biology’, in eLS. John Wiley & Sons, Ltd, pp. 1–7. doi: 10.1002/9780470015902.a0003356.pub2.

10. Dhillon, S. S. et al. (2019) ‘Metabolic profiling of zebrafish embryo development from blastula period to early larval stages’, PLOS ONE. Edited by D. Monleon. Public Library of Science, 14(5), p. e0213661. doi: 10.1371/journal.pone.0213661.

11. Dunn, N. (2018) Raising Larvae in the Zebrafish International Resource Center Autonursery. Available at: https://wiki.zfin.org/display/prot/Raising+Larvae+in+the+Zebrafish+International+Resource+Center+Autonursery (Accessed: 18 June 2018).

12. Giavarina, D. (2015) ‘Understanding Bland Altman analysis Lessons in biostatistics’, Biochemia Medica, 25(2), pp. 141–51. doi: 10.11613/BM.2015.015.

13. Gillis, T. E. et al. (2015) ‘Characterizing the metabolic capacity of the anoxic hagfish heart’, Journal of Experimental Biology. Company of Biologists Ltd, 218(23), pp. 3754–3761. doi: 10.1242/jeb.125070.

14. Howe, K. et al. (2013) ‘The zebrafish reference genome sequence and its relationship to the human genome’, Nature, 496(7446), pp. 498–503. doi: 10.1038/nature12111.

15. Lu, M. J. et al. (2016) ‘Sample Size for Assessing Agreement between Two Methods of Measurement by Bland-Altman Method’, International Journal of Biostatistics. Walter de Gruyter GmbH, 12(2). doi: 10.1515/ijb-2015-0039.

16. Motulsky, H. J. and Brown, R. E. (2006) ‘Detecting outliers when fitting data with nonlinear regression - A new method based on robust nonlinear regression and the false discovery rate’, BMC Bioinformatics. BioMed Central, 7(1), p. 123. doi: 10.1186/1471-2105-7-123.

17. Müller, M. E. et al. (2019) ‘Mitochondrial Toxicity of Selected Micropollutants, Their Mixtures, and Surface Water Samples Measured by the Oxygen Consumption Rate in Cells’, Environmental Toxicology and Chemistry. Wiley Blackwell, 38(5), pp. 1000–1011. doi: 10.1002/etc.4396.

18. Rana, N. et al. (2010) ‘Caffeine-Induced Effects on Heart Rate in Zebrafish Embryos and Possible Mechanisms of Action: An Effective System for Experiments in Chemical Biology’, Zebrafish, 7(1). doi: 10.1089/zeb.2009.0631.

19. Roy, U. et al. (2017) ‘Metabolic profiling of zebrafish (Danio rerio) embryos by NMR spectroscopy reveals multifaceted toxicity of β-methylamino-L-alanine (BMAA)’, Scientific Reports. Nature Research, 7(1), pp. 1–12. doi: 10.1038/s41598-017-17409-8.

20. Santos, L. C. et al. (2017) ‘Caffeine Dose-Response Relationship and Behavioral Screening in Zebrafish’, in The Question of Caffeine. InTech. doi: 10.5772/intechopen.68341.

21. Seth, A., Stemple, D. L. and Barroso, I. (2013) ‘The emerging use of zebrafish to model metabolic disease’, Disease Models & Mechanisms, 6(5), pp. 1080–1088. doi: 10.1242/dmm.011346.

22. Svendsen, M. B. S., Bushnell, P. G. and Steffensen, J. F. (2016) ‘Design and setup of intermittent-flow respirometry system for aquatic organisms’, Journal of Fish Biology. Blackwell Publishing Ltd, 88(1), pp. 26–50. doi: 10.1111/jfb.12797.

23. Svendsen, M. B.S., Bushnell, P. G. and Steffensen, J. F. (2016) ‘Design and setup of intermittent-flow respirometry system for aquatic organisms’, Journal of Fish Biology. Blackwell Publishing Ltd, 88(1), pp. 26–50. doi: 10.1111/jfb.12797.

24. Svendsen, M. B. S., Bushnell, P. G. and Steffensen, J. F. (2019) ‘AquaResp’. doi: http://doi.org/10.5281/zenodo.2584015.

25. Tickle, P. G., Hutchinson, J. R. and Codd, J. R. (2018) ‘Energy allocation and behaviour in the growing broiler chicken’, Scientific Reports. Nature Publishing Group, 8(1), pp. 1–13. doi: 10.1038/s41598-018-22604-2.

26. Zakaria, Z. Z. et al. (2018) ‘Using Zebrafish for Investigating the Molecular Mechanisms of Drug-Induced Cardiotoxicity’, BioMed Research International. Hindawi Limited. doi: 10.1155/2018/1642684.

